# Membrane curvature regulates Ups1 dependent phosphatidic acid transfer across lipid bilayers

**DOI:** 10.1101/2025.04.03.647039

**Authors:** Fereshteh Sadeqi, Dexin Dong, Kai Stroh, Marian Vache, Jutta Metz, Dietmar Riedel, Andreas Janshoff, Herre Jelger Risselada, Caroline Kolenda, Michael Meinecke

**Affiliations:** Heidelberg University Biochemistry Center, Heidelberg 69120, Germany; Department of Physics, Technical University Dortmund, 44221 Dortmund, Germany; Institute for Physical Chemistry, University of Göttingen, 37077 Göttingen, Germany; Department of Structural Dynamics, Max Planck Institute for Multidisciplinary Science, 37077 Göttingen, Germany

**Author notes:** These authors contributed equally to this paper (Fereshteh Sadeqi, Dexin Dong).

## Abstract

Mitochondria are essential organelles in eukaryotic cells, enclosed by two membranes with distinct compositions and functions. In addition to the endoplasmic reticulum, mitochondria are major sites of cellular lipid production. Cardiolipin, for example, is exclusively synthesized in the mitochondrial inner membrane. This requires the precursor lipid phosphatidic acid to be imported from the endoplasmic reticulum to the mitochondrial outer membrane. Subsequently phosphatidic acid is transferred to the inner membrane by the lipid transfer protein Ups1/PRELID1. The regulation of this process, the role of membrane physico-chemical properties, and the mechanisms by which energy barriers are overcome during lipid extraction and insertion remain poorly understood.

Here, we demonstrate that Ups1 exhibits a strong preference for binding to positively curved membrane regions. Our findings reveal that phosphatidic acid extraction is energetically favored at these membrane domains, leading to enhanced lipid transfer between membranes with high positive curvature and we show that events at the donor membrane are rate limiting for the transfer cycle. Our data suggest that Ups1 membrane binding is modulated by pH, lipid composition, and membrane morphology, pointing to a complex, multipartite regulatory network underlying intra-mitochondrial lipid transfer.

## Introduction

Eukaryotic cells exhibit a high degree of compartmentalization characterized by the presence of various membrane-enclosed organelles. On one hand, organellar membranes act as barriers creating specialized reaction spaces. On the other hand, essential biochemical pathways such as lipid biosynthesis and ATP production occur at or within these membranes. Consequently, the composition of organellar membranes is highly diverse in terms of both protein and lipid content and is tailored to their physiological function (van Meer et al., 2008).

The endoplasmic reticulum (ER) is a major site of lipid biosynthesis (Bishop & Bell, 1988). From there, lipids are distributed and transferred to various organelles, including mitochondria. Lipid transfer to mitochondria is non-vesicular and is believed to be mediated by lipid transfer proteins at or near membrane contact sites (Flis & Daum, 2013; Osman et al., 2011; Tatsuta et al., 2014). In mitochondria, some of these lipids are further processed to other lipid species, highlighting the crucial role of mitochondria in cellular lipid biosynthesis. Among these lipids, phosphatidic acid (PA) is converted through a multi-step process in the mitochondrial inner membrane (MIM) to the mitochondrial signature lipid cardiolipin (CL) (Osman et al., 2011). With its two negative charges and strong cone shape, CL plays a crucial role in shaping the biophysical properties of mitochondrial membranes (Konar et al., 2023; Olofsson & Sparr, 2013; Wilson et al., 2019). Beyond this structural role, CL is crucial for many mitochondrial processes such as the functionality of the respiratory chain complexes, the ATP synthase and the protein import machinery, as well as for mitochondrial dynamics, apoptosis, and mitophagy (Dudek, 2017; Gonzalvez et al., 2008; Muhleip et al., 2019; Paradies et al., 2019; van der Laan et al., 2007). Consequently, impairments in CL metabolism result in mitochondrial dysfunction and have been linked to various pathologies, including cardiomyopathies, metabolic disorders, neurodegenerative diseases and cancer. Furthermore, CL levels decline with aging, possibly contributing to increased reactive oxygen species production and a heightened susceptibility to apoptosis (Ahmadpour et al., 2020; Falabella et al., 2021; Paradies et al., 2019; Prola & Pilot-Storck, 2022).

CL biosynthesis in the MIM begins with the precursor lipid PA, which must be imported from the ER and then transported from the mitochondrial outer membrane (MOM) to the MIM. This non-vesicular transport of PA across the intermembrane space (IMS) is mediated by a conserved heterodimeric protein complex found across eukaryotes. In yeast, this complex consists of Ups1 (PRELID1 in mammals), which functions as the lipid-binding protein, and Mdm35 (TRIAP1), which stabilizes Ups1 and protects it from degradation by the i-AAA protease Yme1 and the metallopeptidase Atp23 (Osman et al., 2009; Potting et al., 2013; Potting et al., 2010; Tamura et al., 2009; Tamura et al., 2010).

Crystal structures of the Ups1/Mdm35 complex have revealed that the lipid-binding pocket of Ups1 is formed by a seven-stranded β-sheet which, together with an α-helix on one side, creates a cavity that accommodates a single PA molecule. The PA headgroup is deeply embedded within the pocket, while its acyl chains extend toward the entrance. The acyl chains are shielded from the aqueous environment by the Ω-loop, which is thought to function as a lid (Lu et al., 2020; Miliara et al., 2015; Watanabe et al., 2015; Yu et al., 2015). In vitro studies have shown that Ups1 effectively transfers PA between artificial membrane vesicles (Connerth et al., 2012; Miliara et al., 2015; Watanabe et al., 2015). They also demonstrated that Mdm35 is required to keep Ups1 soluble when not membrane-bound but upon membrane binding of Ups1 dissociates and remains in solution (Connerth et al., 2012).

These findings support the following model: Ups1 binds to the IMS surface of the MOM, likely dissociating from Mdm35. A PA molecule is extracted by Ups1 from the MOM, and Ups1 then reassembles with Mdm35 to form a transferable complex. PA is subsequently shuttled by Ups1/Mdm35 through the aqueous IMS. Finally, Ups1 binds to the IMS surface of the MIM, dissociates from Mdm35, and releases the PA into the MIM (Connerth et al., 2012; Tamura et al., 2020).

The molecular details of many of these steps are poorly understood. It is, for example, unclear what gives directionality to the process, how membrane characteristics regulate Ups1 activity, or how the energy barrier is overcome to break lipid-lipid interactions, extract a lipid from one membrane, and insert it into another. To address this, we combined in vitro and in silico experiments to quantitatively analyze the interaction of Ups1 with membranes and investigate how these interactions regulate lipid transfer.

## Results

### Ups1 exhibits pH-dependent membrane binding with a transition point within the pH range of the IMS

Understanding the membrane-binding properties of Ups1 is crucial for a comprehensive understanding of its lipid transfer mechanism. To analyze the binding characteristics in detail, it is essential to ensure that the interaction occurs via the protein’s natural binding site without interference from additional factors. Ups1 is proposed to interact with negatively charged membranes via positively charged residues (Connerth et al., 2012). In previous studies, constructs containing an N-terminal His-tag were commonly used for purification and biochemical characterization (Connerth et al., 2012; Lu et al., 2020; Watanabe et al., 2015; Yu et al., 2015). Given its proximity to the putative membrane-binding region, the His-tag might potentially affect membrane binding (Figure 1A). With a pKa of approximately 6.0, it introduces additional positive charges in a pH-dependent manner, potentially masking the intrinsic binding properties of Ups1. To address this issue, we constructed and expressed a cleavable Ups1 variant (Supplementary Figure 1B) to eliminate the potential influence of the His-tag and assessed its membrane interactions.

**Figure 1.**
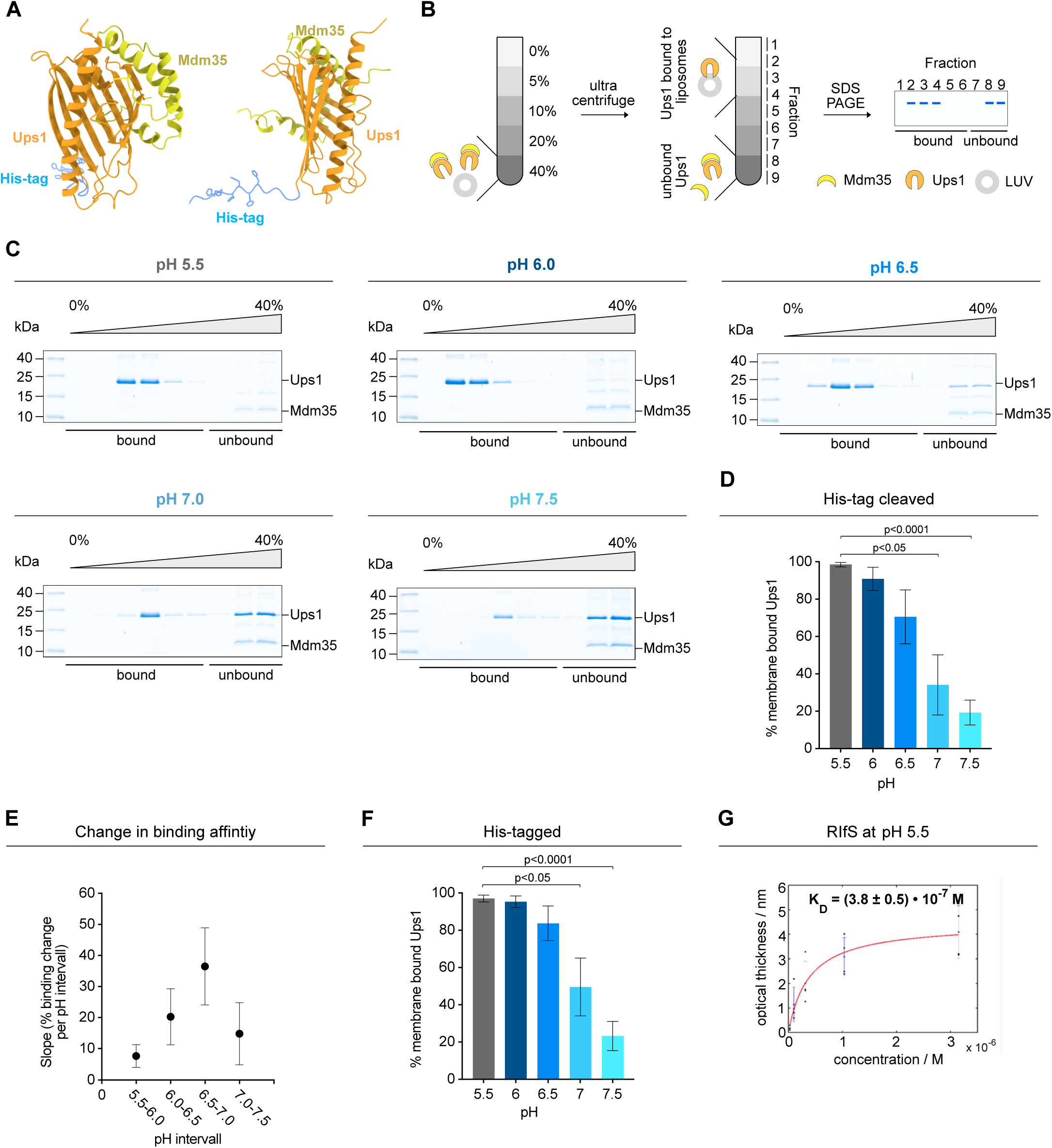
pH-dependent binding of Ups1 to membranes reveals a shift in affinity across the physiological range of the IMS **(A)** Ribbon diagram of His-tagged Ups1 in complex with Mdm35 as predicted by AlphaFold 3 (Abramson et al., 2024). **(B)** Schematic diagram of the flotation assay used to study protein membrane interactions (details see Methods). **(C)** Flotation analyses using untagged Ups1/Mdm35 and LUVs composed of 50% PC, 30% PA, 19.875% PE, 0.125% TF488-PE at indicated pH values. A representative image from three independent experiments is shown for each condition. (**D**) The percentage of membrane-bound Ups1 relative to total Ups1 quantified from Coomassie-stained gels of (C). Data represent the mean ± SD from three independent experiments. Significant differences relative to pH 5.5 are indicated. **(E)** Change in % membrane-bound Ups1 across the indicated pH intervals based on (D). Each data point represents the binding change between indicated pH values calculated using linear regression. Data are shown as mean slope ± SE from three independent experiments. (**F**) Quantification of membrane-bound His1-Ups1/Mdm35 from flotation assays using the same liposomes as in (C) at the indicated pH values. Data represent the mean ± SD from three independent experiments. Significant differences relative to pH 5.5. are indicated. Representative gels are shown in Supplementary Figure 1C. (**G**) Dissociation constant (*K*_D_) of Ups1/Mdm35 binding to membranes was obtained by a Langmuir fit of the optical thickness (red line).

We analyzed membrane binding using flotation assays (Figure 1B). In this assay, proteins are incubated with liposomes to allow binding, then the sample is placed at the bottom of a density gradient and ultracentrifuged. Because liposomes are less dense, they float into the low-density fractions. Ups1 that is membrane-bound coflotates and therefore cofractionates with these liposome-containing fractions. In contrast, unbound Ups1 remains in the higher-density fractions. Using this assay, we observed that at pH 5.5, Ups1 was detected predominantly in the low-density, liposome-containing fractions (identified by TF488-PE fluorescence; Supplementary Figure 5), indicating co-flotation and thus strong membrane binding at low pH. As pH increased, the membrane binding of Ups1 progressively decreased with a transition point near pH 6.5 and relatively low binding at pH 7.5 (Figure 1C-E). No obvious differences between Ups1 and His-Ups1 could be detected (Figure 1F, Supplementary Figure 1C). Reflectometric interference spectroscopy (RIfS) measurements confirmed these findings, yielding a dissociation constant of K_D_ = 0.38 ± 0.05 µM at pH 5.5, while no binding was resolvable at pH 7.4 (Figure 1G).

Our findings reveal a pronounced pH-dependent binding of Ups1 to PA-containing membranes, raising questions about the molecular mechanisms. pH can affect several membrane properties, such as charge, lipid headgroup dipoles, lipid packing, and interfacial tension. These parameters are not controlled in our assays. With regards to membrane charge, the pKa of PA has been reported to be around 6.9 when in proximity to primary amines, such as those found in PE. Based on this consideration, one would expect stronger binding at higher pH due to enhanced electrostatic interactions, a behavior observed with other PA-binding proteins (Young et al., 2010; Zegarlinska et al., 2018; Zhou et al., 2024). However, Ups1 exhibits the opposite trend, indicating that additional factors influence its pH-dependent membrane interactions. Our experiments argue against the involvement of histidines from the His-tag, shifting attention toward intrinsic features of the protein.

pKa predictions indicate that histidines at positions 5, 9, 33, and 100 may have values within the critical pH range, depending on the conformation of Ups1. It is conceivable that protonation events at these sites could induce conformational changes underlying the observed pH-dependent membrane interactions. Clarifying the molecular mechanism underlying the pH-dependent membrane binding of Ups1, as well as its physiological relevance (see discussion), will be an important objective for future investigation.

### Ups1 exhibits curvature-dependent membrane binding, preferentially associating with regions of high positive curvature

The interaction of Ups1 with membranes has been shown to depend on electrostatic interactions, yet its dependency on other membrane characteristics like membrane morphology remains unexplored. Mitochondria exhibit a striking ultrastructure, with their characteristic cristae architecture generating membranes with complex geometries (Barbot & Meinecke, 2016; Sjostrand, 1953; Zick et al., 2009). Beyond this large-scale structural organization, local membrane morphology is influenced by factors such as protein integration, membrane undulations and lipid composition (Graham & Kozlov, 2010; Khalifat et al., 2008; Lee et al., 2025; Tarasenko & Meinecke, 2021). For example, proteins with local curvature-inducing properties have been identified in both the inner and outer mitochondrial membranes, highlighting that local curvature changes are integral features of both compartments (Barbot et al., 2015; Callegari et al., 2025; Davies et al., 2012; Diederichs et al., 2020; Hessenberger et al., 2017; Takeda et al., 2021; Tarasenko et al., 2017). Conversely, membrane curvature plays a crucial role in specific protein-membrane interactions, influencing not only protein binding but also spatial distribution and functional activity (Antonny, 2011; Komatsu et al., 2000; Larsen et al., 2020; Sugiura et al., 2019; Wang et al., 2023).

To investigate whether membrane morphology influences Ups1 binding, we first employed an in silico approach using coarse-grained molecular dynamics (CG-MD) simulations. Curvature within a lipid bilayer system was induced by applying anisotropic compression along the x-axis, allowing expansion along the z-axis. The system was then equilibrated to enable curvature-induced lipid partitioning. Ups1 was introduced into the simulation, and its partitioning free energy (ΔΔF) was determined as a function of membrane curvature (Figure 2A).

**Figure 2.**
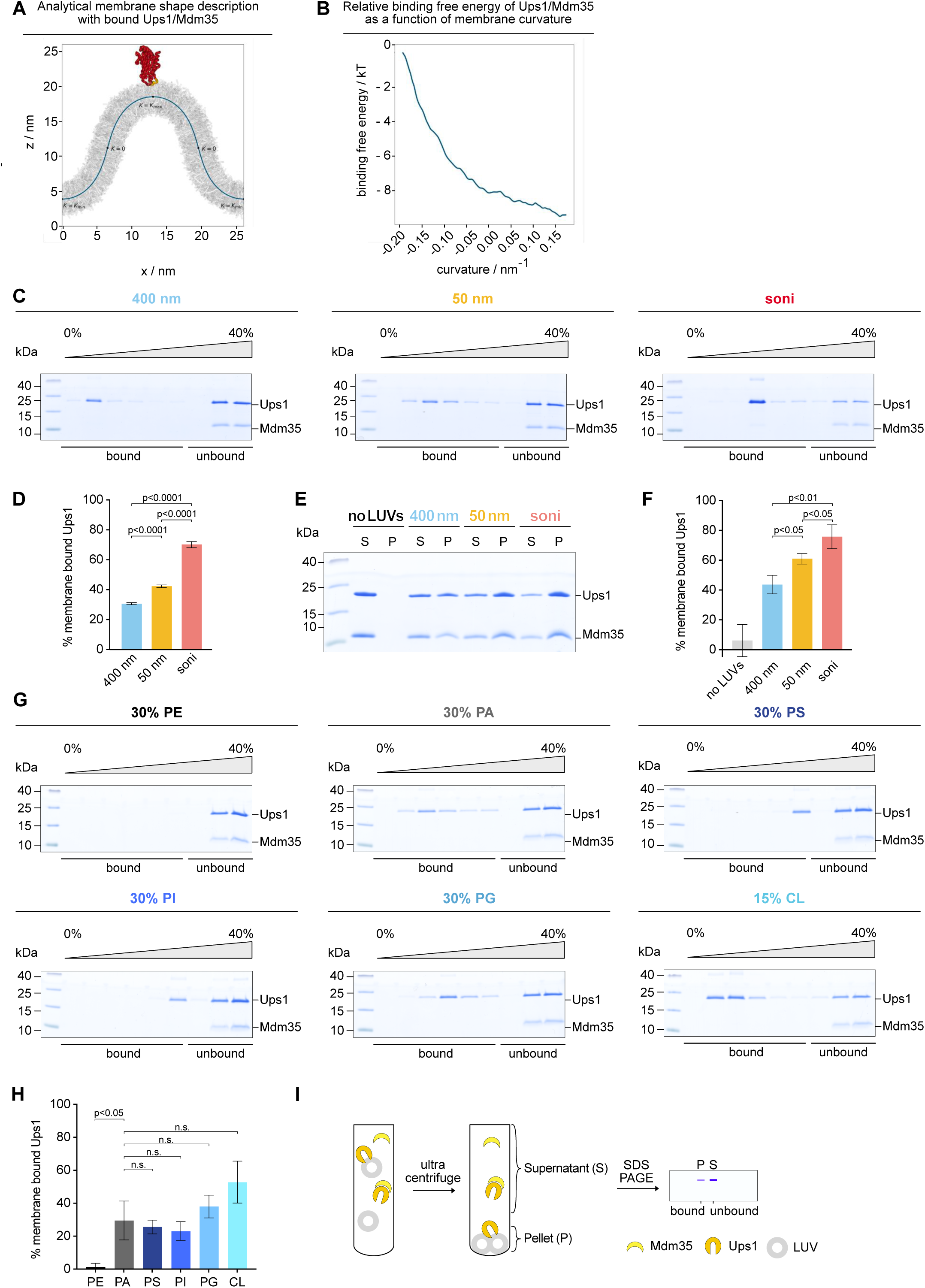
Ups1 membrane binding is dependent on curvature and electrostatic interactions. **(A)** Schematic illustration of the coarse-grained molecular dynamics simulation setup used to investigate the curvature-dependent binding of Ups1 to lipid membranes. The analytical membrane shape description is shown together with the Ups1/Mdm35 complex bound to the membrane. **(B)** Relative binding free energy (ΔΔF) of Ups1/Mdm35 as a function of membrane curvature. **(C)** Flotation analyses showing the binding of untagged Ups1/Mdm35 to liposomes extruded through membranes with indicated pore sizes and composed of 50% PC, 30% PA, 19.95% PE, 0.05% TF488-PE at pH 7.0. A representative image from three independent experiments is shown for each condition. **(D)** The percentage of membrane-bound Ups1 relative to total Ups1 quantified from Coomassie-stained gels of (C). Data represent the mean ± SD from three independent experiments. **(E)** Co-sedimentation analyses showing the binding of untagged Ups1/Mdm35 to liposomes extruded through membranes with the indicated pore sizes. Liposomes were composed of 50% PC, 30% PA, 19.95% PE and 0.05% TF488-PE at pH 7.0. A representative image from three independent experiments is shown for each condition. **(F)** The percentage of membrane-bound Ups1 relative to total Ups1 quantified from Coomassie-stained gels of (E). Data represent the mean ± SD from three independent experiments. **(G)** Flotation analyses showing the binding of untagged Ups1/Mdm35 to liposomes composed of 69.875% PC, 0.125% TF488-PE and 30% of either PE, PA, PS, PI, or PG, as indicated. CL liposomes were composed of 84.875% PC, 0.125% TF488-PE, and 15% CL. Liposomes were extruded through 50 nm membranes and the assay was performed at pH 7.0. A representative image from three independent experiments is shown for each condition. **(H)** The percentage of membrane-bound Ups1 relative to total Ups1 quantified from Coomassie-stained gels of (G). Data represent the mean ± SD from three independent experiments. **(I)** Schematic diagram of the co-sedimentation assay used to study protein membrane interactions (details see Methods).

Across a curvature range from-0.188 nm−1 (K_min_) to +0.188 nm^−1^ (K_max_), we observed a significant change in partitioning free energy of ΔΔF = 9.2 ± 1.5 k_B_T. Thus, the binding free energy decreased with increasing positive curvature, indicating a preferential localization of Ups1 to membrane regions of high positive curvature (Figure 2B). These in silico results suggest that Ups1 has a curvature-sensing capability.

To validate these computational findings, we conducted biochemical assays using liposomes of varying diameters to mimic different degrees of membrane curvature. Liposomes were generated by extrusion through membranes with pore sizes of 50 nm and 400 nm to produce vesicles with high and low positive membrane curvature, respectively. Additionally, 50 nm extruded liposomes were sonicated, resulting in a population that includes vesicles with even smaller diameters. Dynamic light scattering (DLS) measurements confirm the significant size differences (Supplementary Figure 2A). As larger particles scatter more and therefore in DLS even a few larger vesicles can shift the size peak, we performed cryo-EM experiments and analyzed vesicle size distribution (Supplementary Figure 2B). Cryo-EM analysis of 400 nm extruded liposomes shows a broad size distribution with a large fraction of liposomes ranging from 100 to 600 nm, whereas 50 nm extruded liposomes display a peak between 45 and 60 nm, and sonicated liposomes are are mainly in the 15 to 30 nm range. Experiments were performed at pH 7.0, which is within the physiological pH range of the mitochondrial IMS, ensuring that the observed interactions reflect the physiological binding characteristics. Following this experimental setup, liposome co-flotation assays were performed to assess curvature-dependent binding. A relatively weak binding of 31% was observed for 400 nm extruded liposomes, while binding increased to 42% for 50 nm extruded liposomes and reached 70% for sonicated liposomes (Figure 2C&D). In agreement with the flotation analyses, Ups1 preferentially bound to small vesicles in co-sedimentation assays (Figure 2I) with a gradual increase in association as membrane curvature increased (Figure 2E&F). Given that larger liposomes are more prone to multilamellarity, we next tested whether differences in accessible membrane area could account for the apparent curvature preference. Cryo-EM analyses revealed comparable accessible membrane surface area for sonicated and 50 nm liposomes, whereas 400 nm liposomes showed a modest reduction attributable to multilamellarity (Supplementary Figure 3A). However, binding assays across lipid concentrations of 2.5–10 mM showed that especially in otherwise tightly packed membranes, binding to 400 nm liposomes remained low at all concentrations tested. Notably, binding to 400 nm liposomes stayed substantially lower than binding to sonicated liposomes even at fourfold higher lipid concentration (Supplementary Figure 3B). Together, these results support membrane curvature, in addition to negative surface charge, as a key determinant of Ups1 membrane binding.

Thus, our experimental findings align well with the molecular dynamics simulations, collectively demonstrating that Ups1 exhibits a distinct binding preference for positively curved membrane regions. Interestingly, membrane deformation experiments indicate that Ups1 not only senses membrane curvature but also exhibits mild curvature-inducing activity (Supplementary Figure 3C&D). This suggests that, beyond associating with locally curved membrane regions, Ups1 might subsequently stabilize or even induce or further enhance membrane curvature.

### Curvature enhances membrane association of Ups1 but does not independently drive its binding

It has been established that Ups1 binding to membranes depends on the presence of negatively charged lipids in the membrane. This conclusion is based on experiments with large liposomes, focusing on its interaction with relatively flat membranes (Connerth et al., 2012). Our findings demonstrate that, in addition to charge, membrane curvature contributes substantially to Ups1 binding. However, while curvature appears to enhance membrane association, it remains unclear whether it is sufficient to drive binding independently.

To address this, we analyzed Ups1 binding to highly curved neutral membranes and compared its binding to highly curved negatively charged membranes. To ensure comparable surface charge, we aimed to keep the negative charge constant across vesicles containing different negatively charged lipid species, which is especially relevant for CL. Zeta potential measurements confirmed comparable surface charge, showing no significant differences among these vesicles, whereas PE-containing vesicles exhibited a significant shift toward a less negative zeta potential (Supplementary Figure 3E).

Ups1 bound to highly curved negatively charged membranes, with no clear preference for a specific negatively charged lipid species (Figure 2G&H). Compared to PA, CL showed a trend toward stronger binding, however, this effect was not statistically significant under our experimental conditions. Notably, no Ups1 binding was observed to neutral highly curved membranes, indicating that curvature alone is insufficient to support Ups1 binding (Figure 2G&H). Thus, our findings significantly extend previous results obtained using large vesicles at pH 5.5, which demonstrated a strict requirement for negative charges and increased binding to CL-containing membranes (Connerth et al., 2012). This dependency is not only maintained in highly curved membranes but also persists at the physiological pH of 7.0. Taken together, these findings suggest that membrane curvature plays a crucial role in promoting Ups1 binding to membranes and that it acts in concert with electrostatic interactions rather than serving as an independent determinant of membrane association.

### Curvature enhances Ups1 binding to the MOM, while the high negative charge density of the MIM supports strong binding with reduced curvature dependency

The two mitochondrial membranes have a distinct and complex lipid composition. The MIM contains relatively high levels of CL, which contributes through its two negative charges to the high negative charge density of the MIM. In contrast, the MOM contains less CL and also less other negatively charged lipids, resulting in a membrane with less overall surface charge (Zinser et al., 1991). To gain insight into whether curvature sensing is relevant for Ups1 membrane binding in vivo, we assessed the curvature-dependent binding of Ups1 to membranes mimicking the native lipid compositions of both MOM and MIM. Binding to the MOM was significantly influenced by curvature, with 45% of Ups1 binding to flat membranes compared to 69% on highly curved membranes (Figure 3A). This suggests that within the lipid environment of the MOM, Ups1 preferentially associates with membrane regions exhibiting positive membrane curvature. In contrast, binding to the MIM was consistently strong, reaching 86% even on flat membranes (Figure 3B), while a trend toward increased binding to highly curved MIM membranes was maintained. The strong binding of Ups1 to even flat, highly negatively charged membranes suggests that a high density of negative charges can, at least partially, override its curvature-dependent binding, enabling strong association regardless of membrane shape.

**Figure 3.**
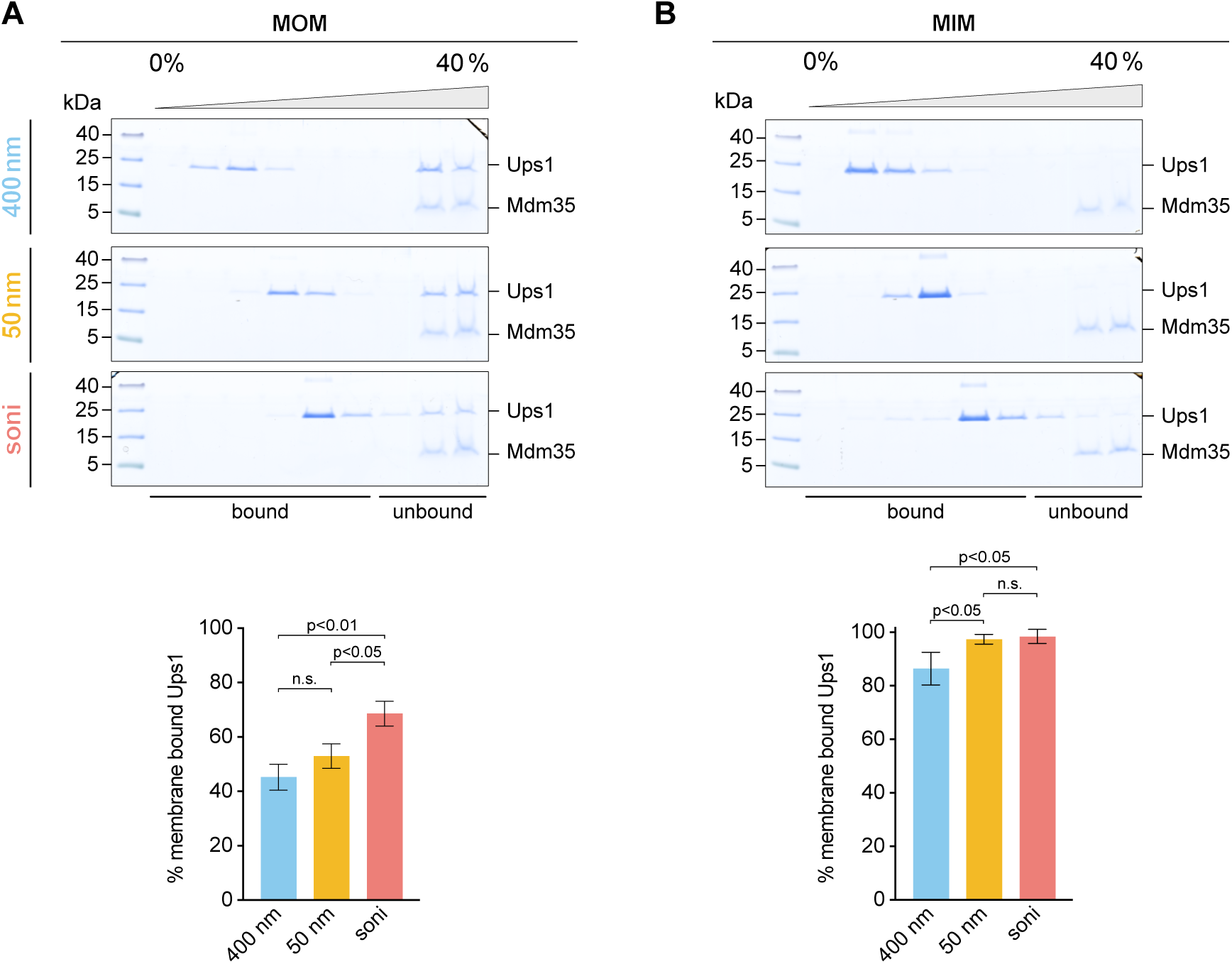
Ups1 binding to membranes mimicking the MOM is curvature-dependent. **(A)** Flotation analyses showing the binding of untagged Ups1/Mdm35 to liposomes extruded through membranes with indicated pore sizes, representing MOM-like membranes (46% PC, 32.95% PE, 10% PI, 1% PS, 6% CL, 4% PA, 0.05% TF488-PE) at pH 7.0. The percentage of membrane-bound Ups1 relative to total Ups1 was quantified from Coomassie-stained gels. Data represent the mean ± SD from three independent experiments. **(B)** Flotation analyses showing the binding of untagged Ups1/Mdm35 to liposomes representing MIM-like membranes (38% PC, 23.95% PE, 16% PI, 4% PS, 16% CL, 2% PA, 0.05% TF488-PE). The experiment and subsequent analysis were performed as in (A).

### Positive curvature increases the free energy of PA, lowering the barrier for lipid extraction

Since we observed a strong curvature-dependent binding of Ups1 to liposomes mimicking the MOM, where lipid extraction occurs, we sought to investigate whether this curvature preference contributes to the efficiency or regulation of the lipid extraction process. Regions of positive membrane curvature, such as the outer leaflet of vesicles, exhibit increased lipid packing defects and surface tension, which enhance the exposure of alkyl chains to solvents (Risselada & Marrink, 2009; Vanni et al., 2014). This energetically elevated state could make lipid extraction more favorable. To test this hypothesis, we employed an in silico approach using coarse-grained molecular dynamics simulations. Using the same buckled membrane previously used for analyzing Ups1 binding, we now focused on lipid sorting behavior (Figure 4A). Curvature-driven lipid partitioning was analyzed, revealing distinct preferences, with PE and PA preferentially localizing to negatively curved regions, while PC accumulated in positively curved regions (Figure 4B). Due to differences in initial concentrations, absolute changes in lipid fractions appear small for PA. However, the relative redistribution of PA along the curvature gradient is substantial, with a decrease of 23% in positively curved regions compared to flat membranes, emphasizing its curvature-dependent sorting behavior. This overall sorting pattern aligns with the properties of the lipid headgroups. Compared to PC, PE and PA have smaller headgroups, making localization of these lipids to positively curved regions less favorable due to a reduced ability to maintain optimal packing. For PA, electrostatic repulsion may also contribute, as its curvature preference is less pronounced compared to PE, despite having an even smaller headgroup.

**Figure 4.**
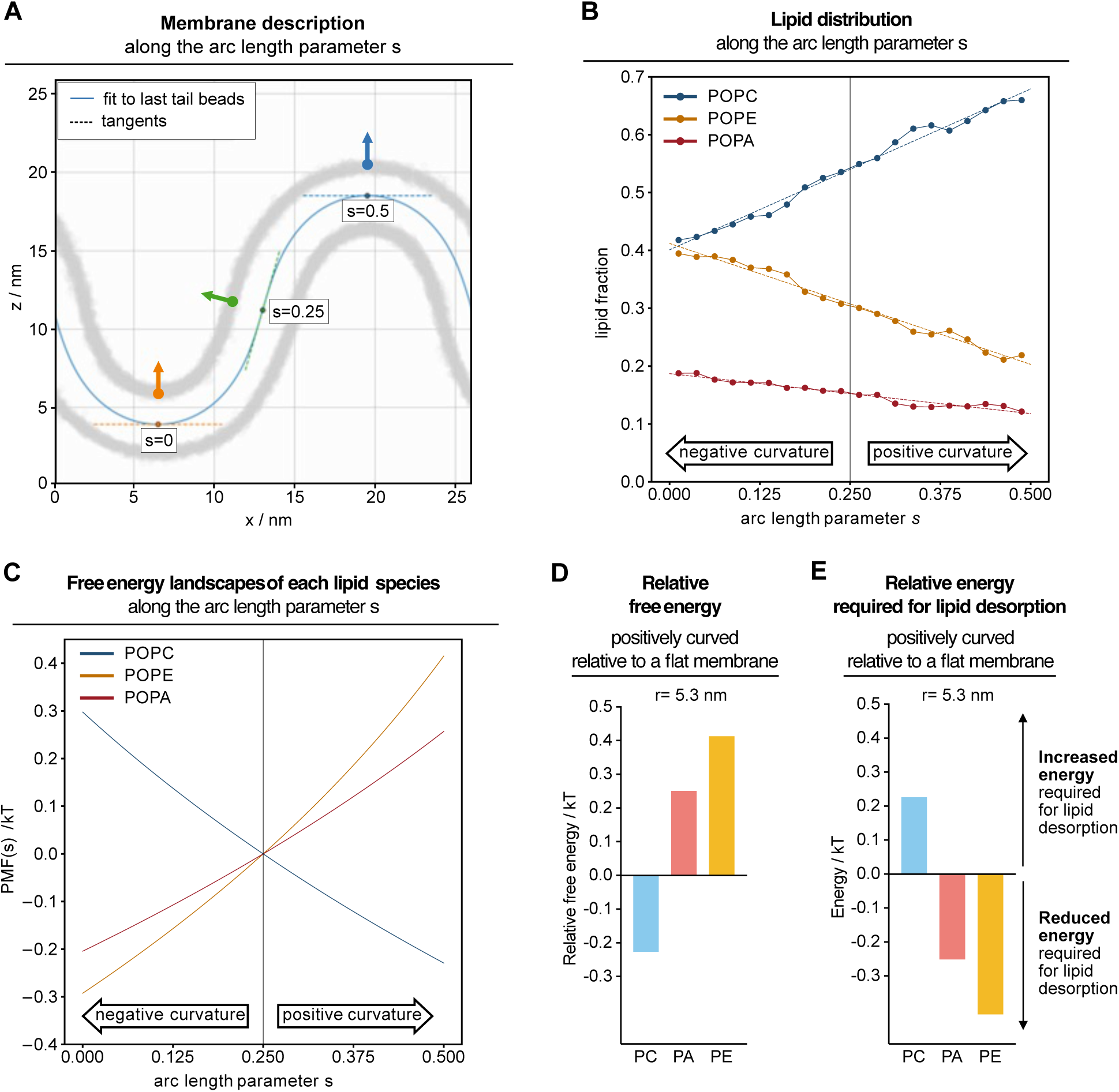
Curvature-dependent lipid partitioning and free energy landscapes reveal energetically favourable PA Extraction from positively curved membranes. **(A)** Schematic illustration of the coarse-grained molecular dynamics simulation setup used to analyse curvature-dependent lipid partitioning. The analytical membrane shape description is shown along the arc length parameter s. **(B)** Lipid distributions along the arc length parameter s after the system was equilibrated to allow curvature-induced lipid partitioning. Errors from bootstrapping are smaller than the symbols. Dashed lines represent the mean of linear fits to every resampled lipid distribution. **(C)** Free energy landscapes of each lipid species along the arc length parameter s constructed from the mean of linear fits to the lipid distributions. **(D)** Relative free energy difference of the indicated lipid species in a positively curved compared to a flat membrane. **(E)** Relative energy differences required for lipid desorption form a positively curved compared to a flat membrane.

To determine the free energy profile of the lipids along the buckled membrane, we constructed the potential of mean force (PMF) from the lipid distributions along the reaction coordinate,

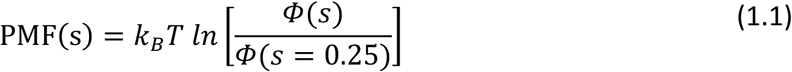

with the Boltzmann constant *k_B_*, the temperature *T*, the arc length parameter *s*, and the lipid distribution φ*(s)*. For PA and PE, the PMF increased from negatively to positively curved membrane regions, indicating that PA and PE are in a higher energy state in positively curved compared to flat membranes. PC followed the opposite trend (Figure 4C&D).

If a lipid is extracted from a region where it is already in a high-energy state, the energy that must be overcome is lower compared to a lipid in a low-energy state. Consequently, because PA has a higher free energy in positively curved regions (Figure 4C&D), the relative energy cost for extraction is reduced in these areas (Figure 4E).

Overall, our in silico results suggest that PA extraction by Ups1 requires less energy from regions of high positive curvature, enhancing the efficiency of lipid extraction and possibly facilitating subsequent transport from these membrane regions.

### Lipid transfer is curvature-dependent, with increased efficiency driven by faster extraction from highly curved membranes

To experimentally test whether membrane curvature has a regulatory effect on Ups1-dependent lipid transfer, we employed a lipid transfer assay (Miliara et al., 2015; Watanabe et al., 2015). In this assay, donor liposomes contain NBD-PA, which is quenched by Rhod-PE in the same membrane. Acceptor liposomes are devoid of any fluorescently labeled lipids. When NBD-PA is transferred from donor to acceptor liposomes, it becomes dequenched, allowing lipid transfer to be monitored as an increase in NBD fluorescence over time (Figure 5A). The initial linear increase in fluorescence reflects the transfer rate. The gradual subsequent slowing of the fluorescence increase indicates that back-transfer becomes relevant, such that not all lipid transfer events contribute to the fluorescence increase.

**Figure 5.**
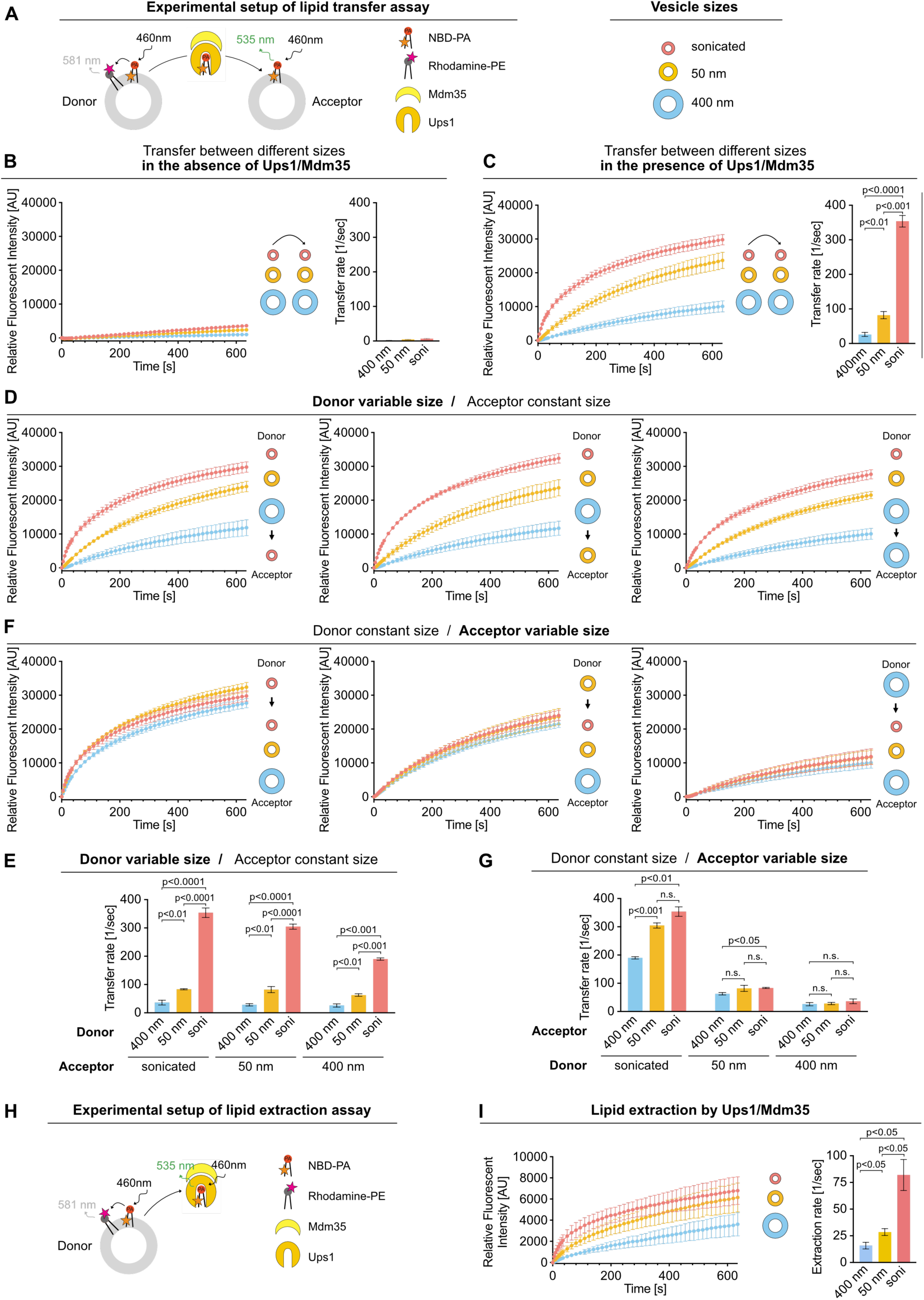
Lipid Transfer by Ups1/Mdm35 Depends on Donor Curvature. **(A)** Schematic diagram of the transfer assay to study Ups/Mdm35 mediated PA transfer. **(B-G)** PA transfer between donor and acceptor liposomes extruded through membranes with the indicated pore sizes. Donor liposomes contained 8% NBD-PA, 2% Rhod-PE, 90% DOPC and acceptor liposomes 10% DOPA and 90% DOPC. **(B)** Size dependent spontaneous PA transfer between donor and acceptor liposomes in the absence of Ups1/Mdm35. Normalized NBD fluorescence is shown over time. Data represent the mean ± SD of three independent replicates (left). The transfer rate is determined from the slope of the linear NBD increase using linear regression (right). Data represent the mean ± SE of three independent replicates. **(C**) Size dependent PA transfer between donor and acceptor liposomes in the presence of Ups1/Mdm35. Same experimental setup and analyses as in (B), with Ups1/Mdm35 added at time point 0. **(D+E**) Effect of donor size on PA transfer by Ups1/Mdm35. Same experimental setup and analyses as in (C). **(F+G)** Effect of acceptor size on PA Transfer by Ups1/Mdm35. Same experimental setup and analyses as in (C). **(H)** Schematic diagram of the extraction assay to study Ups1/Mdm35 mediated PA extraction from donor liposomes. **(I)** PA extraction from donor liposomes with Ups1/Mdm35 added at time point 0. Normalized NBD fluorescence is shown over time. Data represent the mean ± SD of three independent replicates (left). The extraction rate is determined from the slope of the linear NBD increase using linear regression (right). Data represent the mean ± SE of three independent replicates.

To analyze the impact of donor and acceptor size on lipid transfer by Ups1, we compared sonicated, 50 nm, and 400 nm extruded liposomes. Liposomes without the addition of protein resulted in a negligible fluorescence increase, whereas the addition of Ups1 led to a clear increase, indicating successful lipid transfer (Figure 5B&C). Notably, transfer efficiency was strongly influenced by vesicle size (Figure 5C). When both donor and acceptor liposomes were small, transfer rate increased 14-fold compared to relatively flat 400 nm vesicles. The experimental setup was designed to maintain a constant total membrane-binding surface across liposomes of different sizes. It should be noted, however, that this constraint may affect the inter-vesicle spacing during lipid transfer.

To further verify that our results are not driven by the NBD dye attached to the PA molecule, we used a fluorescein dequenching assay that measures the transfer of unlabeled PA. In this assay, fluorescein is dequenched when negatively charged PA molecules are transferred from donor to acceptor liposomes (Supplementary figure 4A) and our results clearly show enhanced transfer rates of non-labelled PA when using smaller liposomes compared to larger ones (Supplementary figure 4B).

To assess the effect of donor size, we performed experiments where the acceptor size was kept constant while the donor size varied (Figure 5D&E). When sonicated acceptor liposomes were used, transfer velocities significantly and gradually increased with higher donor curvature. Similar results were observed for 50 nm and 400 nm extruded acceptor vesicles. This shows that donor size, and thus the curvature of the membrane from which the lipid is extracted, is a strong determinant of transfer rate. Interestingly, acceptor size had a lesser impact in our conditions, as shown by experiments with a constant donor and varying acceptor size (Figure 5F&G). When the donor liposomes were 50 nm or 400 nm, varying the acceptor size had only a minor impact on the transfer rate. When donor liposomes were highly curved (sonicated), a moderate effect of acceptor size on lipid transfer was observed. This indicates that insertion efficiency becomes relevant only when extraction occurs very rapidly, suggesting that extraction is the rate-determining step. Notably, besides extraction, other donor-side processes, such as Ups1 binding or dissociation, could also contribute to defining the transfer rate. Given that the acceptor membrane had a similar composition, one would expect constraints arising from membrane binding and dissociation to occur to a similar extent at the acceptor membrane during lipid insertion. However, this is not what we observe.

To further validate that donor-side processes like lipid extraction are curvature-dependent, we performed an extraction assay (Watanabe et al., 2015). Donor liposomes containing NBD-PA and Rhod-PE were incubated with Ups1 in the absence of acceptor liposomes (Figure 5H). Under these conditions, the fluorescence increase originates exclusively from NBD-PA uptake into Ups1. Consistent with the lipid transfer results, extraction efficiency increased with decreasing donor size (Figure 5I). Extraction was 5-fold faster from sonicated compared to 400 nm liposomes. Taken together, this confirms that lipid extraction is strongly influenced by membrane curvature and is more efficient from highly curved membranes.

## Discussion

In this study, we show that Ups1 membrane binding is regulated by a combination of electrostatic interactions and membrane curvature to optimize lipid transfer efficiency (Figure 6). While electrostatic interactions were previously proposed to mediate Ups1 binding, this was based on experiments conducted at pH 5.5 with His-tagged Ups1 (Connerth et al., 2012), where the positive charge of the His-tag could have influenced binding to negatively charged membranes. We now demonstrate that, even in the absence of the His-tag and at a physiological pH of 7.0, Ups1 binding is dependent on negatively charged lipids, indicating electrostatic interactions with positively charged residues in the protein. Our data further reveal that Ups1 does not exhibit a strict preference for a specific lipid headgroup but rather recognizes membrane charge more broadly. Previous studies suggested a preference for CL, which carries two negative charges (Connerth et al., 2012). These results may have resulted from an overall increase in membrane charge when comparing the same molar amounts of CL with other negatively charged lipids. We maintained a constant total negative charge across lipid compositions by using only half the amount of CL compared to other singly negatively charged lipids. Nevertheless, we observed a trend toward increased binding to CL, though it did not reach statistical significance. Given the proposed role of CL in regulating Ups1 via negative feedback (Acoba et al., 2020; Connerth et al., 2012), the possibility of differential binding to CL compared to other lipids warrants further investigation. However, it should be noted that CL is likely enriched in specific sub-domains within the MIM and might not be readily available as a binding platform at the IMS side of the MIM. Due to its geometry and characteristics of a non-bilayer lipid it will likely sort into regions of negative curvature (Decker & Funai, 2024; Elias-Wolff et al., 2019). Furthermore, it is known to stabilize and, most likely, closely interact with several MIM protein complexes (Duncan et al., 2018; Lange et al., 2001; Musatov & Sedlak, 2017), making it unlikely to be evenly distributed across the plane of the MIM and between both bilayer leaflets. Thus, the extent to which CL regulates Ups1 function remains to be determined.

**Figure 6.**
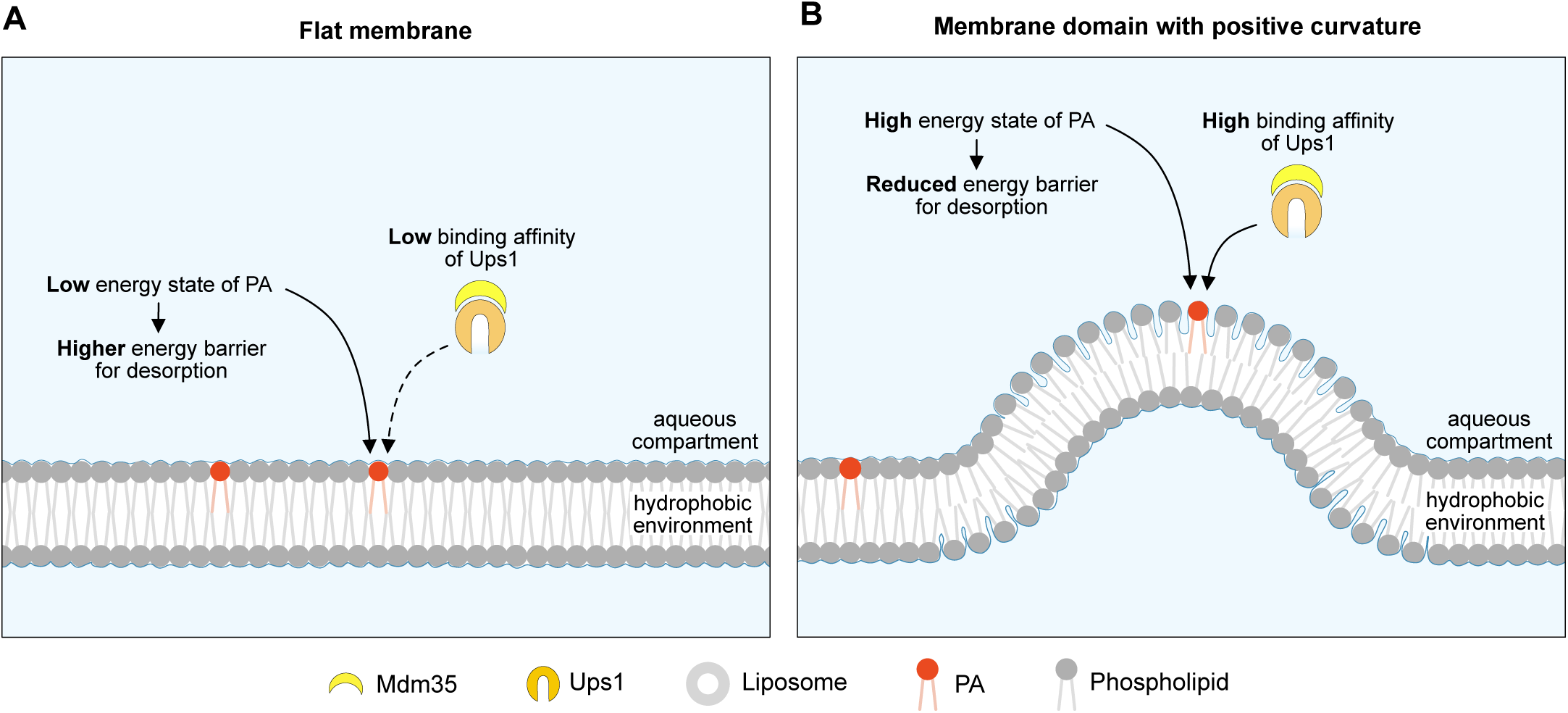
Curvature-Dependent PA Binding and Its Role in Efficient PA Extraction by Ups1 **(A)** Ups1 has low binding affinity to flat membranes, where PA is in a low-energy state, making extraction energetically unfavourable. **(B)** In contrast, Ups1 preferentially binds to highly positively curved membranes, where PA is in a high-energy state, lowering the energy barrier for extraction and enabling more efficient lipid transfer.

Our findings furthermore demonstrate that membrane binding by Ups1 is not solely dictated by electrostatic interactions but is significantly influenced by membrane morphology. Ups1 shows a clear preference for positively curved over flat or negatively curved membranes. However, high curvature alone is insufficient to promote binding, as negatively charged lipids are still required for detectable interaction with highly curved membranes. Interestingly, the dependency on curvature can be largely overridden by highly negatively charged membranes, as observed in our analysis of mitochondrial inner membrane lipid compositions. Under these conditions, Ups1 binding was nearly complete even on flat membranes. Taken together, curvature sensing appears to fine-tune Ups1 binding affinity rather than being the primary determinant of membrane interaction.

For physiological membrane compositions and at the physiological pH of the IMS Ups1 curvature dependent binding was especially apparent for the MOM where PA extraction takes place. Notably, molecular dynamics simulations suggest that PA extraction is energetically more favorable from positively curved membranes. Our lipid transfer experiments are in agreement with this model, as PA extraction was significantly more efficient from highly curved membranes compared to flat membranes. This correlated with a substantial increase in lipid transfer rates. Based on our findings, we propose a model in which Ups1 preferentially binds to positively curved membrane regions where PA exists in an energetically unfavorable state, allowing for efficient extraction. This spatial preference ensures that Ups1 engages with membrane domains where extraction is energetically favorable, thereby increasing lipid transfer efficiency. This mechanism is particularly relevant in environments dominated by flat membranes, such as the MOM, where it helps optimize the use of Ups1 for effective lipid transport. While the MIM contains sub-domains with pronounced curvature, such as cristae junctions, cristae tips, and cristae rims, curvature in the MOM is expected to be less pronounced and more subtle, although such domains are likely to exist. First, we showed that the binding of Ups1 to membranes mildly induces membrane curvature. Thus, protein binding may locally alter membrane morphology exactly at the position where conditions are most favourable for lipid extraction. Furthermore, cryo-EM analyses of outer membrane protein complexes, such as TOM and SAM, have revealed clear signs of membrane distortion in their vicinity (Bausewein et al., 2017; Takeda et al., 2021). These distortions arise from the asymmetrical geometry of the membrane-spanning domains of these proteins. Interestingly, the mitochondrial SAM complex as well as VDAC were recently identified—alongside several other membrane insertases—as lipid scramblases (Jahn et al., 2023; Li et al., 2024), further supporting the notion that the membrane structure in the vicinity of outer mitochondrial membrane proteins is perturbed.

The molecular details by which Ups1 senses membrane curvature remain unclear. Various mechanisms have been described for curvature sensing, including scaffolding and hydrophobic insertion (Has & Das, 2021; Johnson et al., 2024). Scaffolding relies on a binding interface that conforms to membrane curvature, thereby maximizing the contact area between favorable protein-membrane contact sites. This mechanism is highly specific to the geometry of the membrane and the protein. From the known structures of Ups1/PRELID1, though all solved in solution and not in contact with a membrane, it seems unlikely that this mechanism is used. In contrast, hydrophobic insertion is driven by lipid-packing defects induced, for example, by curvature, where amphipathic helices or hydrophobic protrusions insert into the bilayer to stabilize membrane interaction. This mode of binding is not necessarily specific to curvature itself but rather to the packing defects that arise from curvature. Determining which mechanism underlies Ups1 curvature sensing will provide insight into whether Ups1 specifically detects curvature or whether it more generally responds to changes in membrane packing. This distinction is crucial, as it would indicate an additional level of regulation in which Ups1 binding and lipid transfer are influenced not only by membrane charge and curvature but also by other determinants of the membrane that induce packing defects such as acyl chain composition (Vanni et al., 2014).

To gain deeper insight into how Ups1 binds to membranes and senses membrane curvature, it is essential to identify the structural elements responsible for these functions. Curvature sensing is likely an intrinsic property of its membrane-binding interface; however, the lack of structural data for membrane-bound Ups1 makes it challenging to determine the precise molecular features involved. A conformational change upon membrane binding seems likely, as indicated by the fact that Mdm35 must dissociate for Ups1 to efficiently associate with membranes, even though its binding site is distinct from the membrane-binding interfaces predicted by molecular dynamics simulations (Lu et al., 2020; Watanabe et al., 2015). Understanding the conformation of membrane-associated Ups1 will be essential for elucidating the molecular basis of its membrane interactions, including the role of negative charges, potential headgroup specificity—such as a preference for CL—and the mechanisms governing its curvature-dependent association.

Understanding the conformation of membrane-associated Ups1 might also contribute to elucidating the molecular basis of its observed pH-dependent binding to membranes. Since the most significant changes in Ups1 binding with respect to pH align with the physiological pH range of the mitochondrial intermembrane space (IMS), it is an interesting question whether this pH dependency plays a regulatory role in Ups1 function in vivo. Notably, the pH of the IMS fluctuates depending on cellular conditions such as mitochondrial respiration and evidence suggests the presence of a lateral pH gradient (Rieger et al., 2014; Toth et al., 2020). While the presence of a pH gradient across the IMS remains debated, it is plausible that the MIM surface, where protons are pumped by the OXPHOS complexes, could experience a lower pH than the MOM surface, which is thought to be more equilibrated with the cytosol due to the presence of various β-barrel proteins (Benz, 1990). The pH range at which Ups1 undergoes its binding transition suggests that temporal or localized fluctuations in IMS pH could directly modulate Ups1 membrane binding. This dynamic regulation may contribute to establishing directionality or to fine-tune Ups1 activity in response to physiological conditions.

Taken together, our findings demonstrate that the activity of Ups1/Mdm35 is highly sensitive to the lipid composition and morphology of mitochondrial membranes, as well as to pH. Since CL synthesis, a central lipid for mitochondrial function, depends on efficient PA transfer by Ups1/Mdm35, these observations provide insights into how and why alterations in membrane morphology and lipid content may affect CL synthesis and, ultimately, mitochondrial function. Given that Ups1/Mdm35 are structurally and functionally conserved in mammals, these insights may also offer valuable perspectives for understanding the pathophysiology of mitochondrial-related diseases.

## Methods

### Recombinant protein expression and purification

Expression and purification of His-Ups1/Mdm35 was performed as described before with following modifications (Connerth et al., 2012). The coding sequence for *UPS1* carrying an N-terminal His-tag, and *MDM35* from *Saccharomyces cerevisiae* were cloned into the pET-Duet-1 (Merck) vector and co-expressed in Origami B(DE3)pLysS *E.coli* cells (Merck) in the presence of 1 mM IPTG at 30°C for 3h. Cells were harvested and resuspended in lysis buffer (250 mM NaCl, complete protease inhibitor cocktail without EDTA (Roche), 1 mM PMSF, 50 mM Tris/HCl pH 8.0). The lysate was centrifuged at 14,000 rpm for 30 min at 4°C and the supernatant was filtered through a 0.45 µm filter. The clarified lysate was loaded onto a HisTrap FF column (Cytiva). After a wash step with wash buffer (250 mM NaCl, 20 mM imidazole, 50 mM Tris/HCl pH 8.0), proteins were eluted with elution buffer (250 mM NaCl, 500 mM imidazole, 50 mM Tris/HCl pH 8.0) and the protein containing fractions were collected. The protein was further purified by size exclusion chromatography using a HiLoad 16/600 Superdex 75 pg (Cytiva) column with SEC buffer (150mM NaCl, 25mM HEPES pH 7.4). Fractions were analyzed by SDS-PAGE followed by Coomassie blue staining. Selected fractions were pooled and concentrated using an Amicon filter (Merck Millipore).

To obtain Ups1/Mdm35 without a His-tag, a TEV cleavage site was inserted between the His-tag and Ups1 in the previously used pET Duet vector. Expression and purification of His-TEV-Ups1/Mdm35 was performed as described above. The His-tag was removed by incubation with His-tagged TEV protease (van den Berg et al., 2006) in the presence of 3 mM glutathione and 0.3 mM oxidized glutathione. The uncleaved protein and the TEV protease were removed using a HisTrap FF column. Proteins were eluted using an imidazole gradient from 20 mM to 500 mM in 250 mM NaCl, 50 mM Tris/HCl, pH 8.0. Fractions containing the cleaved protein were pooled, rebuffered to 150 mM NaCl, 25 mM HEPES pH 7.4 and concentrated using an Amicon tube.

### Liposome preparation

Liposome preparation was performed as previously described (Denkert et al., 2017). Unless stated otherwise, natural lipids were used as follows: PC (egg L-α-Phosphatidylcholine), PE (egg L-α-Phosphatidylethanolamine), PA (egg L-α-Phosphatidic acid), PS (brain L-α-Phosphatidylserine), PI (liver L-α-Phosphatidylserine) and CL (heart cardiolipin). As indicated, the following lipids were used: DOPC (18:1 (Δ9-Cis) PC), DOPA (18:1 (Δ9-Cis) PA), Rhodamine-PE (18:1 Liss Rhod PE), NBD-PA (18:1-12:0 NBD PA), and TF488-PE (18:1 PE-TopFluor™ AF488).

All lipids were purchased from Avanti Polar Lipids. For liposome formation, lipids were mixed in the desired molar ratios, dried briefly under a stream of nitrogen gas, and dried in a desiccator for at least 2 hours. Dried lipids were hydrated in the respective assay buffer. For the pH-dependent binding assay, assay buffers with the indicated pH were used (150 mM NaCl, 10 mM MES/NaOH, pH 5.5); (150 mM NaCl, 10 mM MES/NaOH, pH 6.0); (150 mM NaCl, 10 mM MES/NaOH, pH 6.5); (150 mM NaCl, 50 mM Tris/HCl, pH 7.0); (150 mM NaCl, 50 mM Tris/HCl, pH 7.5). Unless otherwise indicated, all other experiments were conducted in assay buffer containing 150 mM NaCl and 25 mM HEPES at pH 7.0. Large unilamellar vesicles (LUVs) (vesicles between 100 nm and 1µm) and small unilamellar vesicles (SUVs) (vesicles below 100 nm) were prepared through freeze-thaw cycles, followed by extrusion through polycarbonate membranes (Whatman) with pore sizes of 50 or 400 nm, respectively. To obtain sonicated liposomes, 50 nm extruded liposomes were subjected to sonication (1 min at 50% amplitude and 50% output). For flotation and co-sedimentation assays liposomes were fluorescently labeled with TF488-PE as indicated. The fluorescent intensity of liposomes before and after extrusion was measured and used to determine the liposome concentration after extrusion. For RIfS measurements, the lipids PC/PE/CL/PA were mixed in the following molar ratio 55:20:10:15. After drying, SUVs were generated by swelling the lipid films in buffer (150 mM NaCl, 10 mM MES/NaOH, pH 5.5) followed by sonication.

### Liposome flotation

Flotation assays were performed as previously described (Barbot et al., 2015). 10 µM of the Ups1/Mdm35 complex was incubated with liposomes (5 mM total lipids) in the corresponding assay buffer (see above) for 30 min at 25–30°C in a total reaction volume of 100 µL. The sample was then mixed with Histodenz in assay buffer to a final concentration of 40% Histodenz in a total volume of 800 µL and transferred to the bottom of an ultracentrifuge tube. A discontinuous Histodenz gradient (40%/20%/10%/5%/0%) was created by sequentially overlaying 900 µL each of 20%, 10%, and 5% Histodenz in assay buffer followed by assay buffer on top. The sample was subjected to ultracentrifugation at 45000 rpm (SW60 Ti rotor, Beckman Coulter) for 1 h at the same temperature as above. Nine fractions of 500 µL each were collected, precipitated with 10% TCA, and subjected to SDS-PAGE analysis.

### Co-sedimentation assay

Liposomes were mixed with 10 µM Ups1/Mdm35 in a total reaction volume of 120 µL in the corresponding assay buffer (see above). The sample was incubated for 30 min at room temperature. After incubation the sample was subjected to ultracentrifugation for 30 min at 55000 rpm (TLA-55 rotor, Beckman Coulter) at 25°C. The supernatant was separated from the pellet and both fractions were prepared for SDS-PAGE analysis.

### Tubulation assay

1-2 µl of a lipid solution containing 2 µM lipids of the indicated composition in chloroform was dried on a glass slide. Subsequently, the lipids were rehydrated in 100 µl buffer (50 mM Tris pH 7.0, 150 mM NaCl). Formed lipid membranes were visualized by Differential interference contrast (DIC) microscopy using a CellR imaging station (Olympus) with a x 80 Objective and an ORCA-ER camera (Hamamatsu Photonics) using the Xcellence rt software (Olympus). Images were taken at the indicated timepoints before or after the addition of 10 µM of the indicated proteins and processed using Fiji.

### Reflectometric interference spectroscopy (RIfS) measurements

Reflectometric interference spectroscopy experiments were performed with a home-built setup, consisting of a tungsten halogen light source (LS-1, Ocean Optics, Dunedin, Florida, USA), a flow chamber transparent for visible light which firmly covers a silicon wafer with 5 µm silicon oxide layer (Active Business Company, Brunnthal, Germany) and a spectrometer (Nanocalc-2000-UV/VIS, Ocean Optics). The detailed setup and data analysis to obtain values of the optical thickness is described elsewhere (Stephan et al., 2014).

The flow chamber was first purged with buffer containing 10mM MES, 150 mM NaCL, at pH 5.5. To create a supported membrane, the PC/PE/CL/PA (55:20:10:15) containing SUV dispersion was flushed through the system until reaching a stable plateau of the optical thickness. After rinsing of the supported membrane with buffer, the system was put into a solution of Ups1/Mdm35. In order to obtain a Langmuir isotherm, the experiment was conducted with increasing concentration of Ups1/Mdm35. Lastly, the membrane was rinsed with buffer. To analyse the RIfS data, the obtained optical thicknesses were plotted against the concentration and a Langmuir isotherm was fitted to the data according to eq. 1.

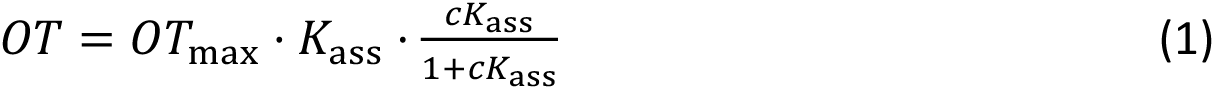

With *OT* being the optical thickness, *OT*_max_ the equilibrium optical thickness at the maximum concentration of protein, *c* the protein concentration and *K*_ass_ the thermodynamic association constant. The thermodynamic dissociation constant (*K*_d_) is the inverse of the association constant.

### Electron Microscopy of LUVs

LUVs in the presence or absence of protein were bound to a glow discharged carbon foil covered 400 mesh grid and stained with 1% uranyl acetate solution as described before (Vasic et al., 2020). Electron microscopic imaging and visualization of the sample was performed at room temperature with a Talos L120C transmission electron microscope (ThermoFischer Scientific, Eindhoven, and The Netherlands).

### In silico curvature sensing

Coarse-grained molecular dynamics simulations were performed with GROMACS 2020.3 (Abraham et al., 2015) using the Martini 2.2 force field (de Jong et al., 2013; Monticelli et al., 2008). The atomistic protein structure (PDB 4XHR, (Yu et al., 2015)) was translated to a coarse-grained description using the martinize Python script (de Jong et al., 2013). Protein secondary structure was determined employing DSSP (Kabsch & Sander, 1983; Touw et al., 2015) and overall protein structure was stabilized via elastic band with a force constant of 500 kJ/(mol*nm^2^).

Initial configurations of the protein membrane system were generated with the python script insane (Wassenaar et al., 2015). 674 lipids per leaflet (371 POPC, 202 POPE, 101 POPA) were put in the x-y plane of a 40 nm * 10 nm * 20 nm simulation box. The protein complex was placed approximately 1 nm above the upper leaflet. The system was solvated with standard Martini water and a 0.15 M NaCl concentration. Steepest-descent energy minimization and initial equilibration (10 ns NVT and 50 ns NPT) were followed by compressing the system in x-direction by applying a pressure of 3 bar in x-direction while allowing the system to expand in z-direction only (Berendsen barostat (τ_p_=12.0 ps, compressibility of 3 * 10^-4^ bar^-1^)(Berendsen et al., 1984)). From the compression trajectory a frame close to the chosen compression level was selected. The selected frame was then used as a starting configuration of a long equilibration phase that was 5 microseconds long (Parrinello-Rahman barostat (τ_p_=12.0 ps, compressibility of 3 * 10^-4^ bar^-1^)(Parrinello & Rahman, 1981)). This long run is necessary to allow curvature-induced lipid partitioning to finalize. For the umbrella sampling the buckled membrane was fixed to its analytical shape by position restraints on the headgroups of lower leaflet lipids with a small force constant (10 kJ mol^-1^ nm^-2^). Influence on upper leaflet dynamics is minimal with these restraints. To generate starting configurations for the umbrella sampling, the protein complex was pulled along the membrane in x-direction. 60 windows equidistant in arc length parameter s, between s = 0 and s = 0.5 were used. Umbrella sampling runs were 1.05 µs long, with the first 0.05 µs for equilibration. Harmonic potentials with a force constant k = 100 kJ mol^-1^ nm^-2^ were used to restraint the hydrophobic loop to a defined position along the reaction coordinate. Umbrella sampling simulations were performed in the NVT ensemble. All simulations were coupled to a constant heat bath using the rate rescale algorithm (Bussi et al., 2007).

Lipid concentrations along the arc length parameter s were extracted from a 2 µs simulation in the NVT ensemble. System setup was as described above, except the protein complex was not present in this simulation. Since the membrane shape is mirror symmetric around s = 0.5, lipid distributions are averaged over both sides and presented from s = 0 to s = 0.5. Potentials of mean force were constructed from linear fits to the lipid distributions. Errors were estimated via bootstrapping (Newman & Barkema, 1999). Each lipid distribution was resampled 1000 times.

### Lipid transfer assay using NBD-PA/Rhodamine-PE

NBD-PA transfer was analyzed using a fluorescence dequenching assay as described (Miliara et al., 2015; Watanabe et al., 2015) with the following modifications. 3.125 µM donor liposomes (90% DOPC, 2% Rhodamine-PE, 8% NBD-PA) and 12.5 µM acceptor liposomes (90% DOPC, 10% DOPA) were incubated with 120 nM Ups1 at 25 °C. NBD fluorescence was recorded over time using a Pherastar BMG Labtech plate reader. The fluorescence intensity measured 0.6 s after injection was set to 0 for normalization (relative fluorescent intensity). The transfer rate was determined by linear regression analysis of the initial linear NBD increase.

### Fluorescein-based lipid transfer assay

Lipid transfer of unlabeled PA was monitored using a fluorescein-based transfer assay adapted from Richens et al. (Richens et al., 2017). Donor liposomes (90% DOPC, 10% DOPA) and acceptor liposomes (100% DOPC) were prepared as described above in assay buffer (25 mM HEPES pH 7.0, 150 mM NaCl). Donor liposomes were labeled by adding fluorescein–DHPE in ethanol (final ethanol ≤0.1% v/v) and incubating for 1.5 h at 37 °C in the dark. Free fluorescein–DHPE was removed using a PD-10 column (G-25 MiniTrap, Cytiva) equilibrated in assay buffer. Reactions contained 100 µM donor, 400 µM acceptor, and 3 µM Ups1. Fluorescein fluorescence was recorded over time using a PHERAstar plate reader (BMG Labtech). The fluorescence intensity measured 0.6 s after injection was set to 0 for normalization (relative fluorescence intensity).

### Dynamic Light Scattering

DLS measurements were performed with the Dynapro Nanostar (Wyatt Technology). Liposomes were diluted to 1mM with the appropriate buffer for the measurement. The temperature was set to 25°C and 20 DLS acquisitions were taken for each sample. The DYNALS algorithm was used for the regularisation fit.

### Zeta potential measurements of liposomes

The zeta potential of liposomes was measured with the ZetaSizer Nano Z instrument and DTS1070 cuvettes (Malvern Panalytical). Liposomes were diluted to a concentration of around 1mM for the measurements. The temperature was set to 25°C, dispersant refractive index to 1.332, the viscosity to 0.9021 and dispersant dielectric constant to 78.5. The particle refractive index was set to 1.42. The data was processed with the Smoluchowski model with F(𝜅a) set to 1.5. Due to the salt content in the samples, the voltage was set at 10V. For each sample, six measurements with 40 sub-runs were taken. DLS measurements were also performed as described above to confirm the liposomes were of the same size.

### Cryo-EM

A cryogenic transmission electron microscope (cryo-TEM) equipped with an autoloader (Glacios, ThermoFisher Scientific, Waltham, USA) enabled visualization of the morphology and size of 400 nm-, 50 nm-extruded or sonicated liposomes liposomes. A volume of 3 µL with a concentration of around 10mM of the dispersion was added to a glow-discharged Quantifoil (R 2/1, copper) specimen support grid, blotted in a humidified atmosphere for 5 sec (Vitrobot, ThermoFisher Scientific, Waltham, USA) operated at 6°C, and immediately plunged into liquid ethane cooled with liquid nitrogen. Afterward, the autogrids were stored under liquid nitrogen. The samples were examined with the electron microscope at 200 kV and images were taken at a nominal magnification of 57,000 (calibrated pixel size of 2.5 Å) with a Falcon 3 camera operated in linear mode.

The diameters of 263 400 nm-extruded liposomes, 765 50nm-extruded liposomes and 735 sonicated liposomes were measured using FIJI. In addition, multilamellar liposomes were counted and the diameter of all enclosed layers measured to estimate the fraction of accessible membrane.

### pKa predictions

pKa values were predicted using PROPKA version 2.0 via the VEGA online server (Li et al., 2005; Pedretti et al., 2021) based on PDB structures 4XHR, 4YTW, and 4YTX (Watanabe et al., 2015; Yu et al., 2015).

### Statistical analyses

Significance was assessed using a two-tailed t-test. If the F-test to compare variances detected a significant difference, Welch’s correction was applied. A significance threshold of *p* < 0.05 was used.

### Use of AI tools

AlphaFold 3 (Abramson et al., 2024) was used to generate protein structural models. ChatGPT (OpenAI) was utilized for formulation refinement in the writing of this manuscript.

## Acknowledgments

We thank Dr. Rainer Beck for help with the tubulation assay. Furthermore, we would like to acknowledge access to the cryo-EM infrastructure and support provided by the Cryo-EM Network at the Heidelberg University (HDcryoNet), which is funded and supported by the German Research Foundation (DFG), the Federal Ministry of Education and Research (BMBF) and the Ministry of Science Baden-Württemberg, among others, within the framework of the Excellence Strategy of the Federal and State Governments of Germany. The authors are grateful for the computational resources provided by the Jülich Supercomputing Center and the HLRN Berlin / Göttingen. Furthermore, the authors gratefully acknowledge the data storage service SDS@hd supported by the Ministry of Science, Research and the Arts Baden-Württemberg (MWK) and the German Research Foundation (DFG) through grant INST 35/1503-1 FUGG.

This work was funded by the Deutsche Forschungsgemeinschaft (DFG, German Research Foundation) FOR2848/2 (Project-ID 401510699) and SFB1638/1 (Project-ID 511488495).

---

**Supplementary Figure 1.**
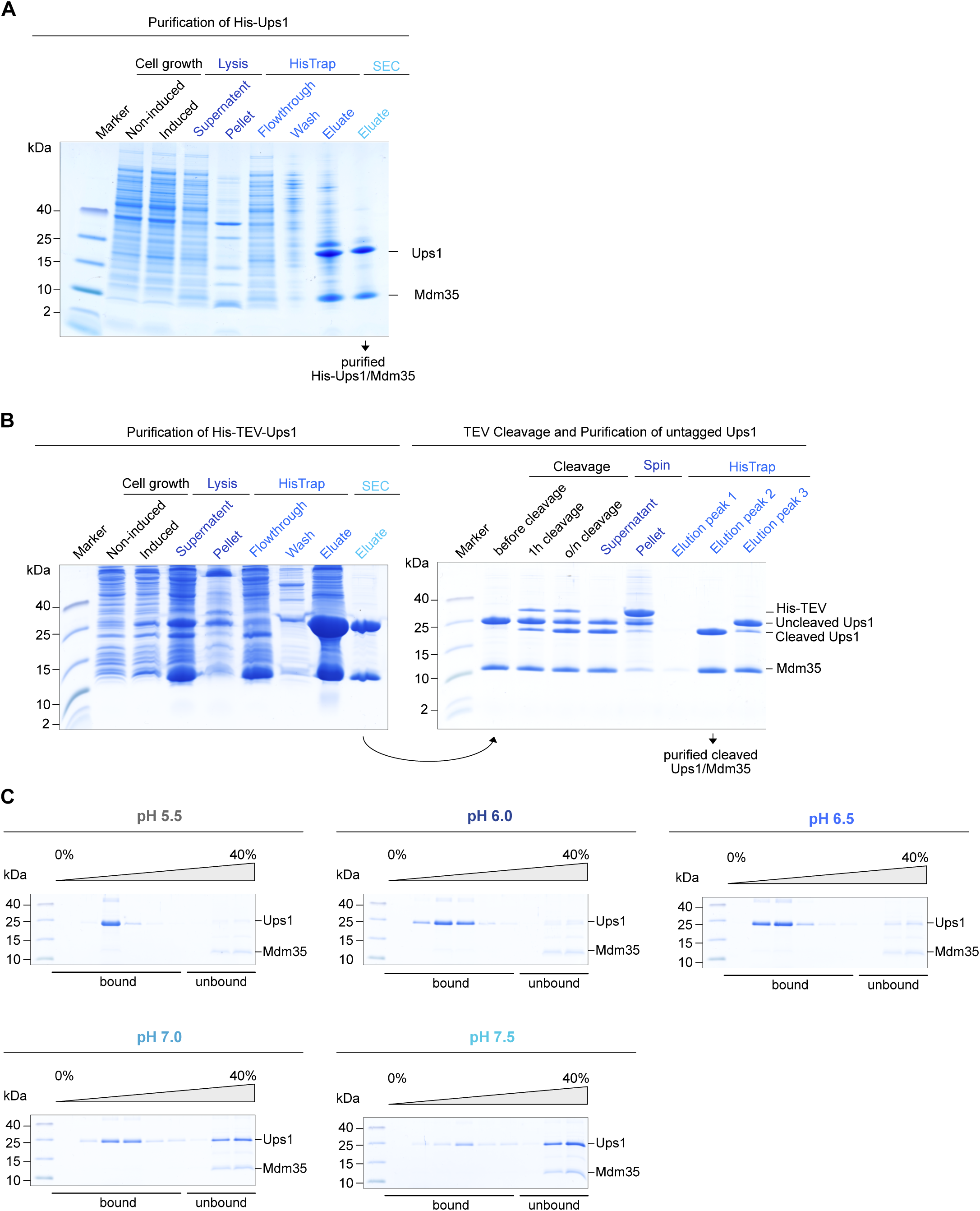
Purification of Ups/Mdm35 and pH dependent binding of His-Ups1/Mdm35. (**A**) Expression and purification of recombinant His-Ups1/Mdm35 complex analyzed by SDS-PAGE and colloidal Coomassie staining. **(B)** Expression and purification of recombinant His-TEV-Ups1/Mdm35, followed by TEV cleavage and subsequent purification of cleaved Ups1/Mdm35, analysed by SDS-PAGE and colloidal Coomassie staining. **(C)** Flotation analyses using His-Ups1/Mdm35 and LUVs composed of 50% PC, 30% PA, 19.875% PE, 0.125% TF488-PE at indicated pH values. A representative image from three independent experiments is shown for each condition.

**Supplementary Figure 2.**
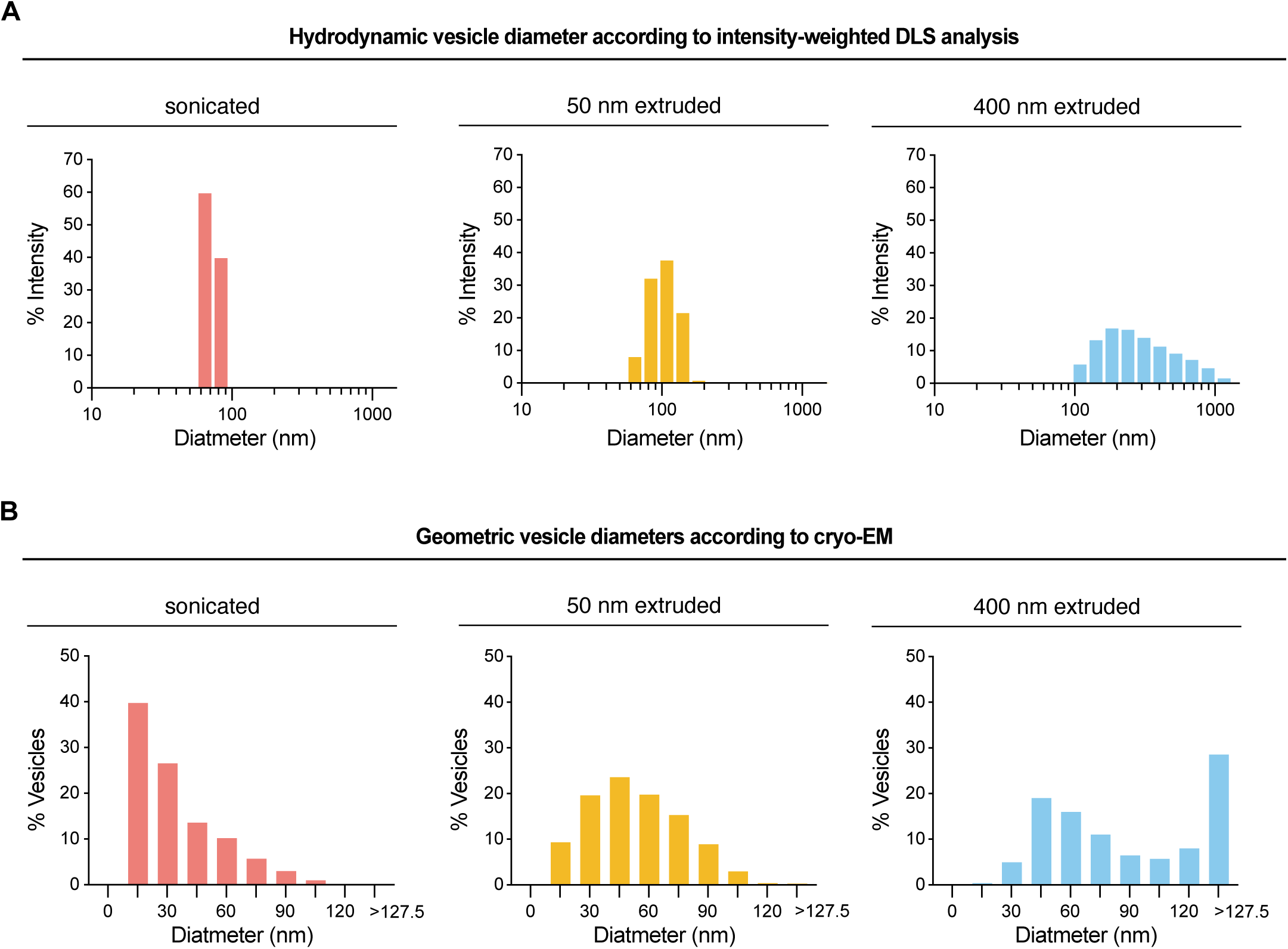
Liposome size distributions measured by DLS and cryo-EM. **(A)** Hydrodynamic vesicle diameter distributions for liposomes extruded through 400 nm or 50 nm membranes or prepared by sonication were determined by intensity-weighted dynamic light scattering (DLS). For each sample, 20 acquisitions were recorded, and the hydrodynamic diameters were obtained using the DYNALS regularization algorithm. **(B)** Geometric vesicle diameter distributions were determined by cryo-EM for liposomes extruded through 400 nm or 50 nm membranes or prepared by sonication. Vesicle diameters were measured in FIJI (n = 263 for 400 nm–extruded, n = 765 for 50 nm–extruded, n = 735 for sonicated).

**Supplementary Figure 3.**
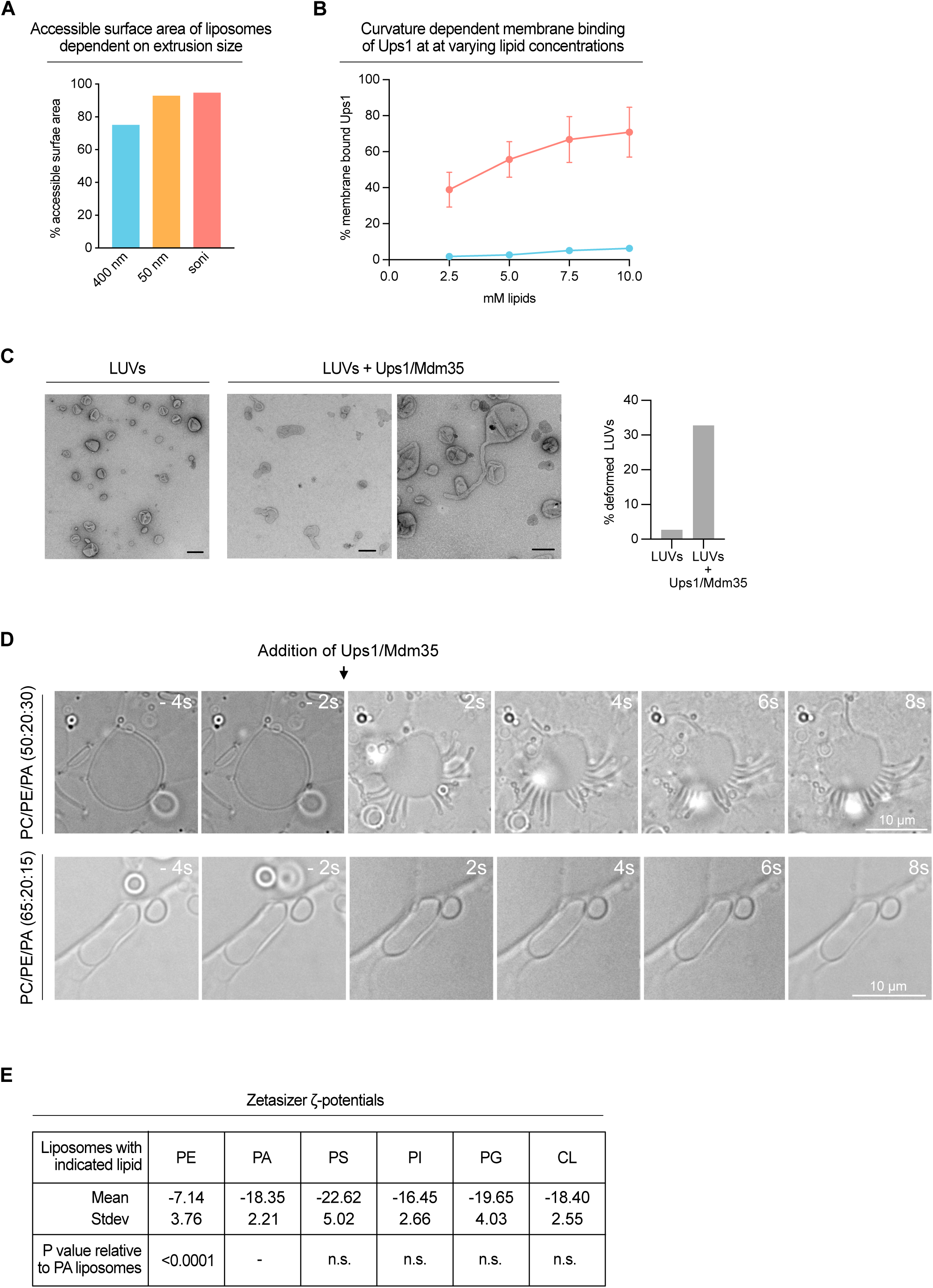
Ups1 shows curvature-and electrostatics-dependent membrane binding and exhibits mild curvature-inducing activity. **(A)** Cryo-EM–based quantification of accessible liposome surface area. Cryo-EM images were used to measure vesicle diameters and, for multilamellar vesicles, the diameters of the outer and all enclosed membrane layers. The fraction of accessible membrane was then quantified from these measurements and is shown as % accessible surface area for 400 nm extruded, 50 nm extruded, and sonicated liposomes. **(B)** Flotation analyses showing binding of untagged Ups1/Mdm35 to liposomes extruded through membranes with the indicated pore sizes over a range of total lipid concentrations. Liposomes were composed of 79.875 mol% POPC, 20 mol% POPA, and 0.125 mol% 18:1 TopFluor AF488-PE (pH 7.0). The percentage of membrane-bound Ups1 relative to total Ups1 was quantified from Coomassie-stained gels. Data represent the mean ± SD from three independent experiments. **(C)** Electron micrographs of liposomes (LUVs) in the absence and presence of Ups1/Mdm35 and quantification of spherical and deformed shapes in control LUVs and in LUVs after incubation with Ups1/Mdm35. A minimum of 200 membranous structures were counted per condition. Scale bar corresponds to 100 nm. **(D)** Tubulation assay in the absence and presence of Ups1/Mdm35. Indicated lipid compositions were dried on a glass slide and rehydrated using assay buffer (50 mM Tris pH 7.0; 150 mM NaCl). Formed lipid membranes were visualized by DIC imaging at the indicated timepoints before or after the addition of 10 µM Ups1/Mdm35. **(E)** Zeta potentials of liposomes prepared with the lipid compositions used for the experiments in Fig. 2G– H.Liposomes contained 69.875% PC, 0.125% TF488-PE, and 30% of the indicated lipid (PE, PA, PS, PI, or PG). CL liposomes contained 84.875% PC, 0.125% TF488-PE, and 15% CL. ζ-potentials were measured by electrophoretic light scattering (Zetasizer). Values are reported as mean ± SD. P values are shown relative to PA-containing liposomes (n.s., not significant).

**Supplementary Figure 4.**
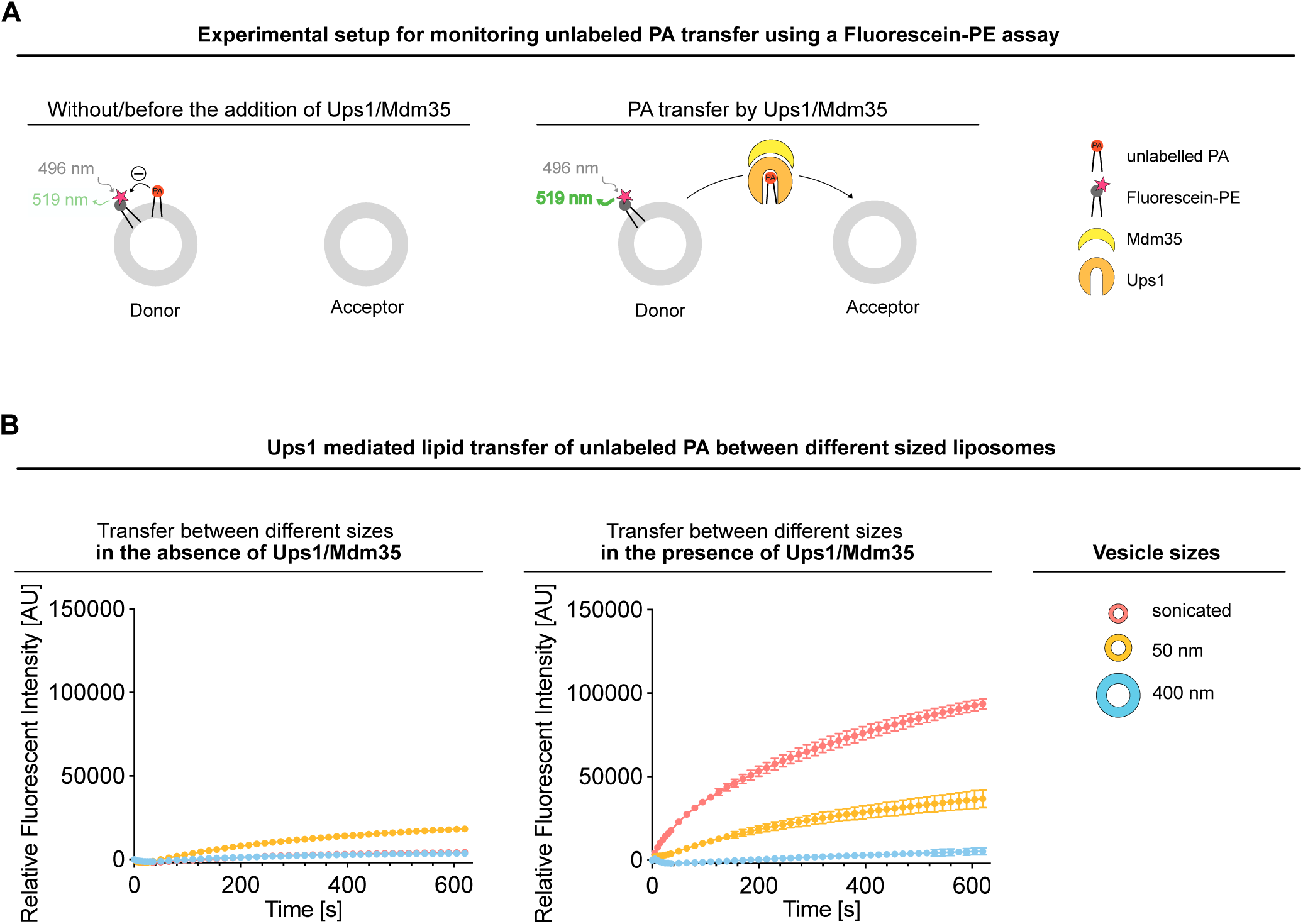
Transfer of unlabeled PA by Ups1/Mdm35 is curvature-dependent, as measured by a fluorescein-based lipid transfer assay. **(A)** Schematic diagram of the fluorescein-based transfer assay to study Ups1/Mdm35-mediated transfer of unlabeled PA. Donor liposomes contain fluorescein–PE and PA, while acceptor liposomes are unlabeled. Fluorescein fluorescence intensity depends on membrane surface potential and thus on membrane surface charge, with increased negative surface charge reducing fluorescence; during PA transfer from fluorescein-labelled donor to acceptor liposomes, the donor membrane becomes less negatively charged and fluorescein fluorescence increases. **(B)** Size-dependent transfer of unlabeled PA between donor and acceptor liposomes in the absence (left) or presence (right) of Ups1/Mdm35. Donor liposomes (90% DOPC, 10% DOPA) were labeled with fluorescein–PE and mixed with acceptor liposomes (100% DOPC). Both vesicle populations were prepared at the indicated sizes (sonicated, 50 nm extruded, or 400 nm extruded). Fluorescein fluorescence was recorded over time and normalized by setting the intensity 0.6 s after injection of Ups1/Mdm35 to zero (relative fluorescence intensity). Data represent the mean ± SD of three independent replicates.

**Supplementary Figure 5.**
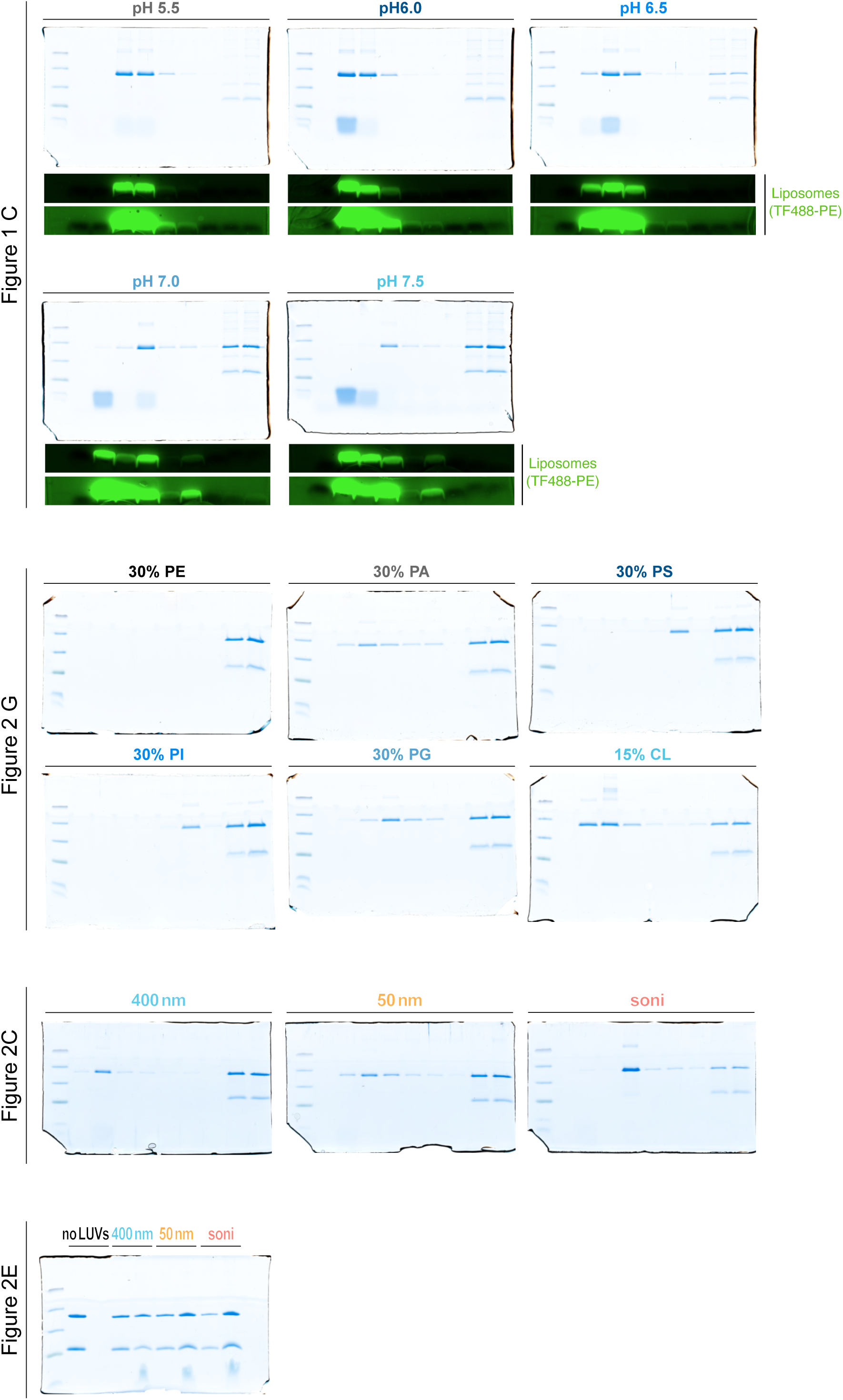
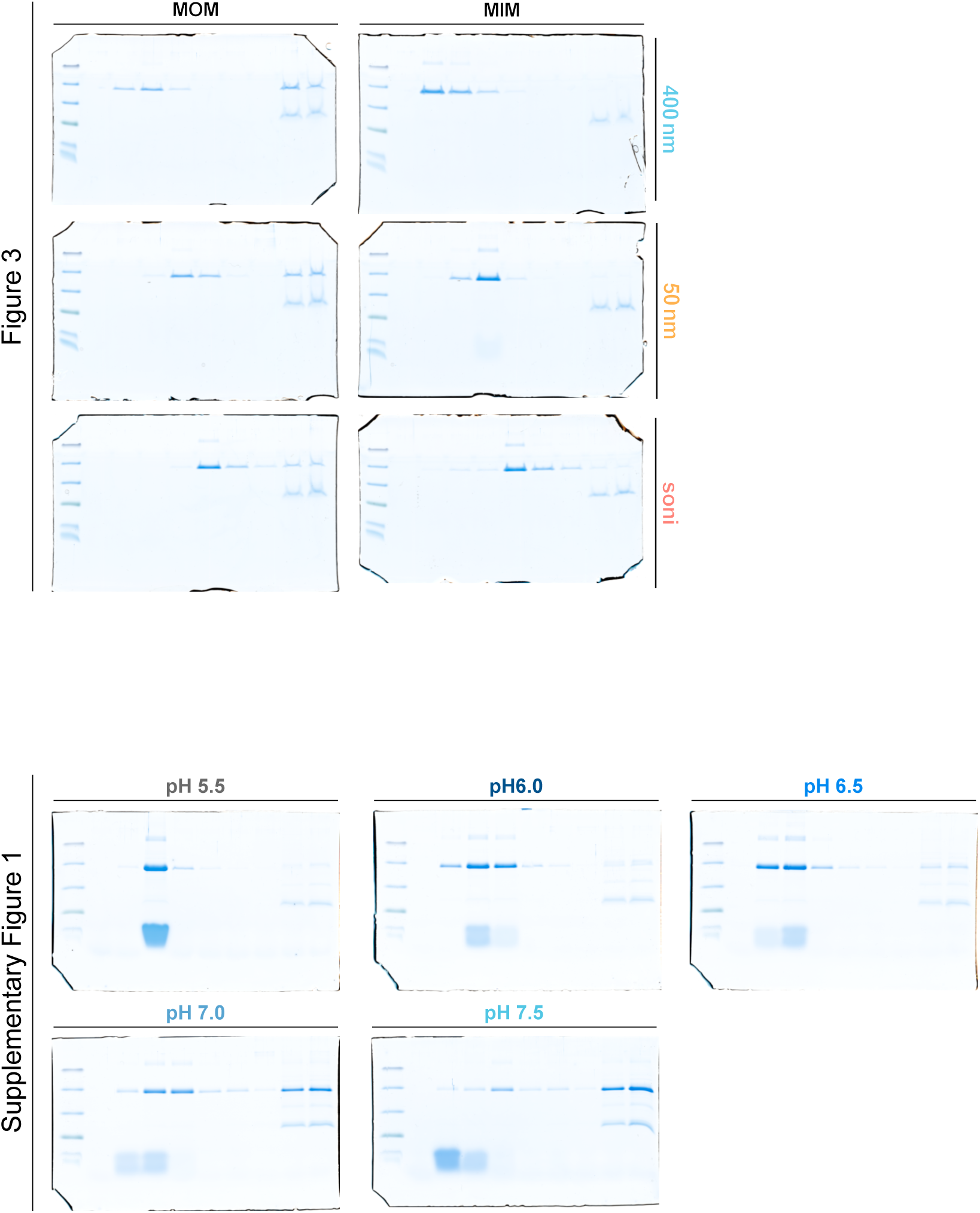
Uncropped SDS–PAGE gels and corresponding TF488-PE fluorescence Images of uncropped SDS–PAGE gels corresponding to the cropped gel images shown in Figures 1–3 and Supplementary Figure 1. For the flotation assays in Fig. 1B, TF488-PE fluorescence was imaged and is shown in addition to indicate the distribution of lipids across fractions. Two contrast and brightness adjustments are shown to visualize both high and low lipid amounts in the different fractions. Ups1 in the low-density fractions cofractionated with liposomes across all conditions shown, indicating co-flotation. Notably, at higher pH Ups1 predominantly cofloated with the subset of liposomes recovered towards intermediate density fractions, consistent with preferential association of Ups1 with vesicles exhibiting higher buoyant density (e.g. due to higher protein loading or vesicle remodeling or clustering).

